# Targeting TRIP13 in Wilms Tumor with Nuclear Export Inhibitors

**DOI:** 10.1101/2022.02.23.481521

**Authors:** Karuna Mittal, Benjamin P. Lee, Garrett W. Cooper, Jenny Shim, Hunter C. Jonus, Won Jun Kim, Mihir Doshi, Diego Almanza, Bryan D. Kynnap, Amanda L. Christie, Xiaoping Yang, Glenn S. Cowley, Brittaney A. Leeper, Christopher L. Morton, Bhakti Dwivedi, Taylor Lawrence, Manali Rupji, Paula Keskula, Stephanie Meyer, Catherine M. Clinton, Manoj Bhasin, Brian D. Crompton, Yuen-Yi Tseng, Jesse S. Boehm, Keith L. Ligon, David E. Root, Andrew J. Murphy, David M. Weinstock, Prafulla C. Gokhale, Jennifer M. Spangle, Miguel N. Rivera, Elizabeth A. Mullen, Kimberly Stegmaier, Kelly C. Goldsmith, William C. Hahn, Andrew L. Hong

## Abstract

Wilms tumor (WT) is the most common renal malignancy of childhood. Despite improvements in the overall survival, relapse occurs in ~15% of patients with favorable histology WT (FHWT). Half of these patients will succumb to their disease. Identifying novel targeted therapies in a systematic manner remains challenging in part due to the lack of faithful preclinical *in vitro* models. We established ten short-term patient-derived WT cell lines and characterized these models using low-coverage whole genome sequencing, whole exome sequencing and RNA-sequencing, which demonstrated that these ex-vivo models faithfully recapitulate WT biology. We then performed targeted RNAi and CRISPR-Cas9 loss-of-function screens and identified the nuclear export genes (*XPO1* and *KPNB1*) as strong vulnerabilities. We observed that these models are sensitive to nuclear export inhibition using the FDA approved therapeutic agent, selinexor (KPT-330). Selinexor treatment of FHWT suppressed *TRIP1*3 expression, which was required for survival. We further identified *in vitro* and *in vivo* synergy between selinexor and doxorubicin, a chemotherapy used in high risk FHWT. Taken together, we identified XPO1 inhibition with selinexor as a potential therapeutic option to treat FHWTs and in combination with doxorubicin, leads to durable remissions *in vivo*.

## INTRODUCTION

Wilms tumor (WT) is the most common childhood renal tumor and represents ~6% of all pediatric cancers with a peak age of presentation at 3 years^1–3^. In the United States, African-American children have 2.5 times higher rates of WT when compared to Caucasians or Asian-American children^4–6^. For low-risk disease, current therapy includes the use of surgery and chemotherapy (e.g., vincristine, dactinomycin). For high-risk disease (e.g. pulmonary metastasis or invasion of the renal sinus and capsule), doxorubicin and radiation therapy are added to low-risk disease therapy. Despite significant increases in response and survival over the past 50 years, ~15% of patients with WT recur^7–9^ and salvage regimens are successful only in 50% of patients and carry significant morbidity.

A major limiting factor in testing novel targeted therapies in WT is the lack of faithful *in vitro* preclinical models. Prior cell line models of WT have been recharacterized as other pediatric cancers such as rhabdoid tumor (e.g. G401) and Ewing sarcoma (e.g. SK-NEP)^10–13^. Recent efforts to generate WT cell lines have been limited due to finite passaging^14–17^. Despite this limitation, a repository of WT organoids and patient derived xenografts (PDXs) has been developed in recent years^18–20^.

Nomination of rational therapeutics for organoid or *in vivo* PDX studies, however, requires systematic *in vitro* efforts using faithful cancer cell lines. Here, we have developed faithful cell line models using genome and transcriptome sequencing of WT which recapitulate known WT biology. We then performed functional genomic screens focused on druggable targets to nominate WT therapeutics^21–23^.

## RESULTS

### WT cell lines faithfully recapitulate genomic and transcriptomic features of WT

Recent studies have shown the feasibility of generating short-term WT cell lines with limited genomic testing ^14–17^. We developed 10 short-term WT cell lines from 8 patients (8 patient derived cell lines and 2 PDX derived cell lines following one passage of the PDX in mice (Annotated as T2); **Fig 1a**; **Methods**). Six patient samples were obtained at time of diagnosis and two were obtained at time of recurrence (Aflac_2377T and CCLF_PEDS_0002_T). Seven patients had favorable histology Wilms tumor (FHWT) and one patient had diffuse anaplastic Wilms Tumor (DAWT; CCLF_PEDS_0023_T).

**Figure 1:**
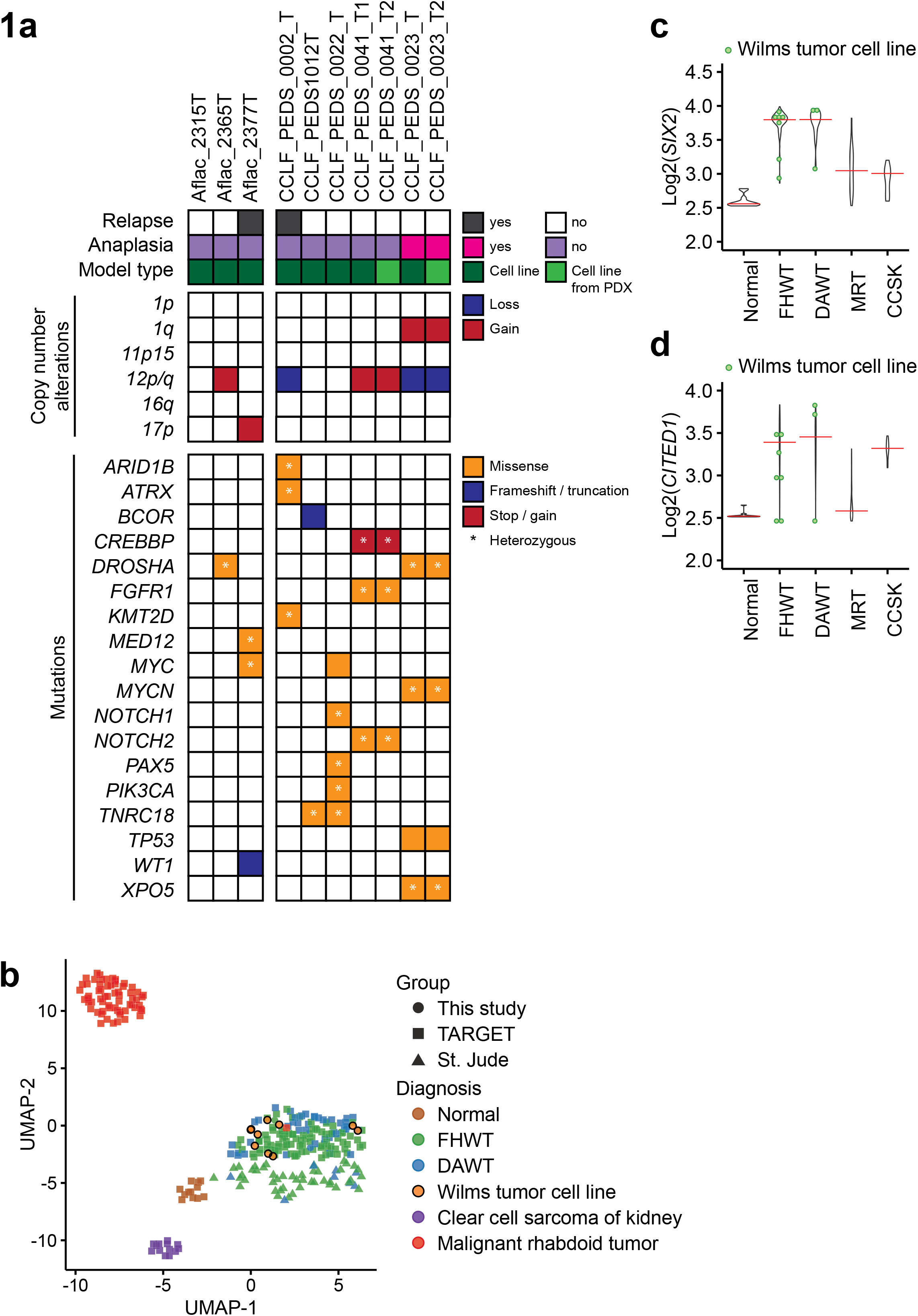
Overview of the Genomic and transcriptomic analysis of Wilms tumor cells lines: **(a)** Co-mutation plot representing the clinicopathological information (top panel), copy number alterations (middle panel), and mutations (bottom panel) in the WT cell lines. T or T1 represents the cell line derived from the patient while T2 represents the cell line derived from the first passage PDX. **(b)** Two-dimensional representation of RNA-seq data using uniform manifold approximation and projection (UMAP) demonstrates high concordance between TARGET (squares) and St. Jude primary (triangles) WT (green - FHWT and blue - DAWT) and our Wilms tumor cell lines (orange circles with black boundaries). Normal tissues samples, clear cell sarcoma of the kidney, and malignant rhabdoid tumor all were clustered separately. **(c)** and **(d)** Dot plots representing the higher expression of commonly dysregulated genes (*SIX2 and CITED1)* in WT in the TARGET and St. Jude datasets and WT cell lines. Green dots indicate WT cell lines.

We then performed ultra-low coverage whole genome sequencing (WGS) to infer copy number status, whole exome sequencing (WES) to identify known mutations in WT and RNA-sequencing to assess the transcriptome in the patients’ tumor and the matched cell lines. We identified 1q gain, a poor prognostic factor in Wilms tumor biology, in one patient (CCLF_PEDS_0023_T) ^24,25^. We observed gain in chromosome 12 in two patients but did not identify LOH in 1p or 11p15. We then assessed the mutational profiles of these tumors and cell lines (**Fig 1a**; **Supp Table 1**). We found that the mutations reflected the spectrum of mutations seen in the WT samples profiled in the NCI Therapeutically Applicable Research to Generate Effective Treatments^26^ (TARGET; **Fig 1a**). Finally, we identified *TP53* (c.469G>T) in CCLF_PEDS_0023_T which is consistent with the pathological diagnosis of anaplastic Wilms tumor ^26^ (**Fig 1a**).

Next, we created a two-dimensional map of the tumor and cell line transcriptomes using the uniform manifold approximation and projection (UMAP) ^27^ and compared it to TARGET^28^ and St. Jude Children’s Research Hospital’s WT datasets ^19^ (**Methods**). We observed that our WT cell lines clustered closely with the WT tumor samples (**Fig 1b)**. We further observed that *SIX2* and *CITED1*, genes involved in WT biology, were upregulated in our WT cell lines consistent with WT biology (**Fig 1c and d**)^29–31^. Collectively, our findings suggest that our patient-derived cell lines recapitulate known biology and serve as faithful representations of WT.

### RNAi and CRISPR-Cas9 screens identify XPO1 as a potential therapeutic target in WT

We then asked if we could identify genetic vulnerabilities in WT despite the short-term lifespan of these cell lines. Following rapid expansion of these cell lines within the first five passages from obtaining the patient’s sample, we subjected a subset of cell lines to targeted loss of function RNA interference and CRISPR-Cas9 screens. Through these screens, we identified genes that lead to decreased cell viability when suppressed or deleted. As cell numbers were limited, we used the Druggable Cancer Targets (DCT) libraries, a library of 429 genes, as previously described ^23,32^ (**Fig 2a**).

**Figure 2:**
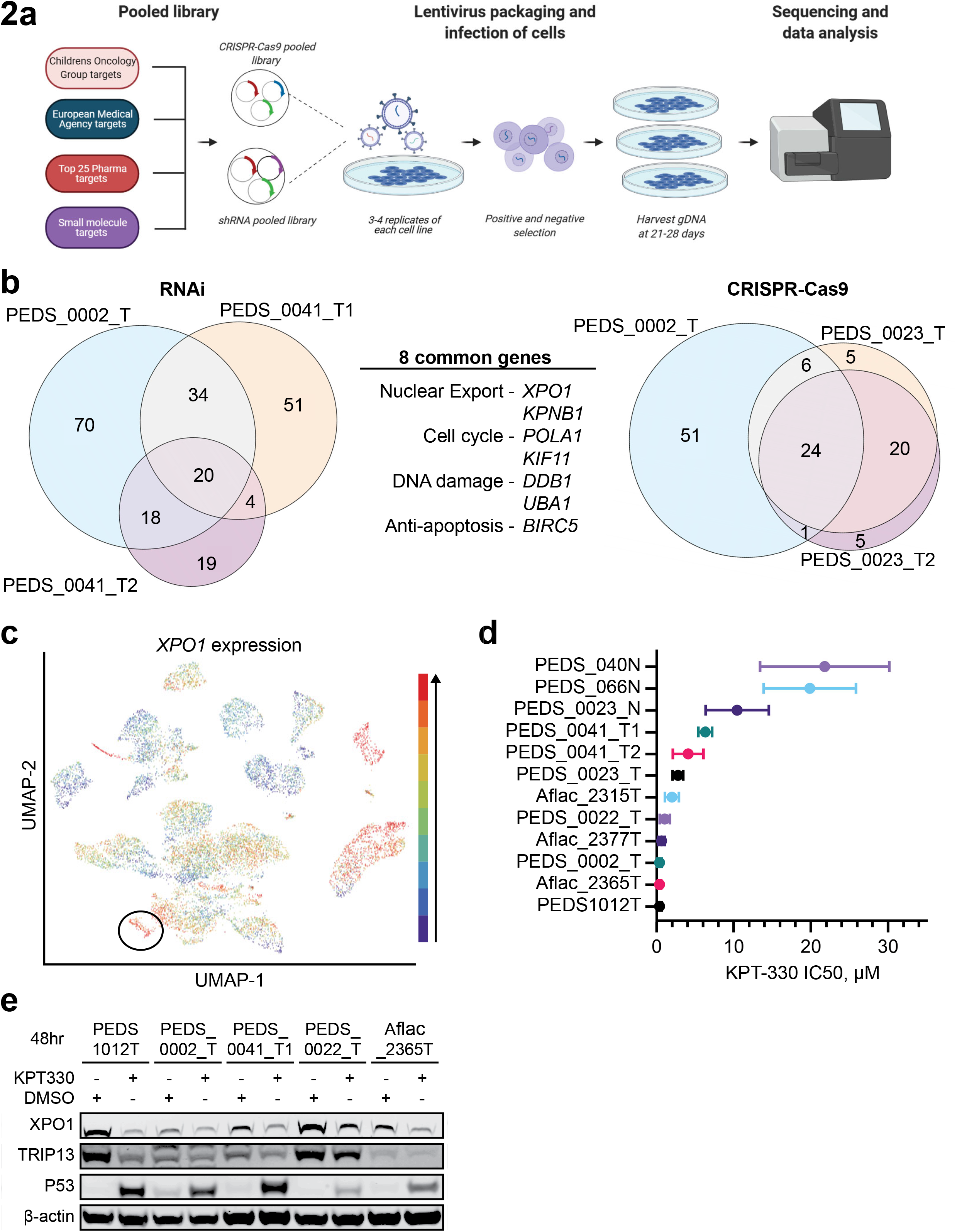
XPO1 is a potential therapeutic target in Wilms tumor cells. **(a)** Schematic outlining the methodology of CRISPR-Cas9 and RNAi functional screens**. (b)** RNAi suppression in three cell lines identified 20 common genes which were critical for the survival of WT cells. CRISPR-Cas9 screens identified 24 common genes in three cell lines which were critical for the survival of WT cells. Eight genes overlapped between the RNAi and CRISPR-Cas9 loss-of-function screens. These 8 genes can be categorized under their role in nuclear export, cell cycle, DNA damage and apoptosis. **(c)** UCSC Treehouse transcriptional data from 12,719 samples showing expression of *XPO1* in all tumor types with WT samples circled in black. Blue to red colors signify expression levels with red being the highest among this cohort. **(d)** Forest plot representing the IC_50_ of KPT-330 in the panel of WT cell lines and normal cells (ending with N). Cell viability data is based on biological duplicates with technical triplicates. **(e)** Immunoblots depicting the decrease in protein levels of XPO1 upon treatment with KPT-330. Data shown are representative of at least biological replicates.

For the RNAi screens, we introduced the DCT lentiviral library into the CCLF_PEDS_0002_T, CCLF_PEDS_0041_T1, and CCLF_PEDS_0041_T2 cells and compared the abundance of each shRNA at the time of infection to its abundance after 25-29 days of cell culture (**Supp Fig 1a–c**). We then used Model-based Analysis of Genome-wide CRISPR/Cas9 Knockout (MAGeCK) to identify 20 genes required for survival across these cell lines ^33^ (**Fig 2b**). We subsequently performed CRISPR-Cas9 screens in CCLF_PEDS_0002_T, CCLF_PEDS_0023_T and CCLF_PEDS_0023_T2 to identify 24 genes that when deleted led to decreased viability across these cell lines (**Supp Fig 1d–f**). From these orthogonal screens, we identified eight genes which overlapped. These included genes involved in nuclear export (*KPNB1* and *XPO1*), regulators of the cell cycle (*KIF11* and *POLA1*), DNA damage (*UBA1* and *DDB1*) and cell survival (*BIRC5*) (**Fig 2b).**

We focused on the role of nuclear export since inhibitors such as selinexor (KPT-330) have been developed and recently FDA-approved in multiple myeloma and diffuse large B-cell lymphoma ^34^. We first assessed the expression levels of *XPO1* across 85 cancer types through the University of California Santa Cruz Treehouse Childhood Cancer Initiative ^35^. From 12,719 patient samples, we found 85.4% of Wilms Tumor samples were in the top 20% of samples with high *XPO1* levels (**Fig 2c**). To validate the requirement of XPO1 in WT cell lines, we assessed cell viability of WT cells using KPT-330 and XPO1 suppression with RNAi (**Fig 2d, Sup fig 2a–b**). Specifically, we determined the IC_50_ values for our WT cell lines and compared these to adjacent normal kidney cell lines (PEDS_0023_N, PEDS_066N, PEDS_040N). We observed that cell lines from adjacent normal renal tissue were not sensitive to KPT-330 with an IC_50_ ranging 10-20 uM (**Fig 2d, Supp fig 2c**). In contrast, our WT cell lines had up to 10-fold lower IC_50_ consistent with responses seen in multiple myeloma where KPT-330 has FDA approval^36^. Specifically, half of these cell lines showed an IC_50_ at nanomolar concentrations (e.g. 25-800nM) whereas the other cell lines had an IC_50_ in the low micromolar range (e.g., 1-5 µM). On-target activity of KPT-330 requires transcriptional upregulation of *XPO1* due to an auto-feedback loop in conjunction with suppression of XPO1 protein levels ^37^. However the downstream mechanisms of action varies by cancer type. Consistent with these prior findings, we saw upregulation of *XPO1* by qRT-PCR and down regulation of XPO1 protein in WT (**Fig 2e and Supp Fig 2d)**. Taken together, these findings suggest that XPO1 is a potential therapeutic target in WT.

### XPO1 inhibition induces cell death through the TRIP13/ TP53 axis

We next investigated a potential mechanism of action of the nuclear export inhibitor KPT-330 in WT. KPT-330 is known to affect tumor suppressors such as a TP53 through nuclear accumulation of TP53 and prevention of MDM2/4 related degradation^38^. We focused our efforts in understanding the mechanisms in FHWT. We treated TP53 wild-type CCLF_PEDS_0041_T1 with DMSO or KPT-330 and performed RNA sequencing (**Methods**). We then performed differential expression analyses and then examined gene sets enriched or suppressed upon KPT-330 treatment using Gene Set Enrichment Analyses (GSEA)^39^. We found 412 genes differentially expressed (**Fig 3a**) and 50 hallmark gene sets significantly enriched or suppressed (**Supp Table 3**).

**Figure 3:**
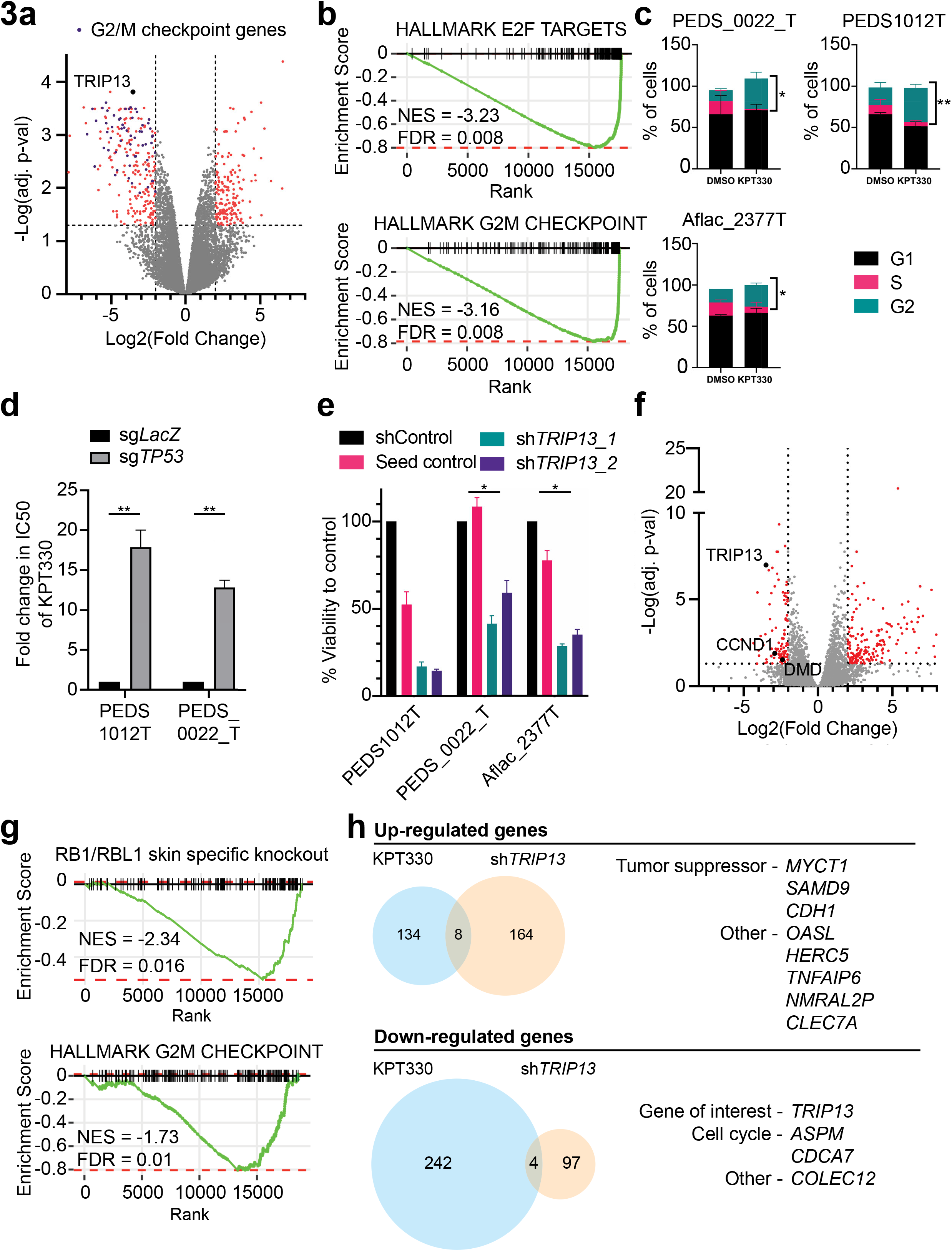
XPO1 inhibition leads to decreased viability through TRIP13 and TP53 axis: **(a)** Volcano plot representing differential gene expression between the KPT-330 and DMSO treated CCLF_PEDS_0041_T cell line. Scattered points represent genes: the x-axis is the fold change for KPT-330 vs DMSO treated CCLF_PEDS_0041_T cells as compared to the p-values. Purple dots represent genes in the Hallmark G2/M Checkpoint gene set. **(b)** Gene set enrichment analysis (GSEA) enrichment score curves for the E2F and G2M hallmark pathways in the CCLF_PEDS_0041_T cells treated with KPT-330. The green curve denotes the ES (enrichment score) curve, the running sum of the weighted enrichment score in GSEA. **(c)** KPT-330 and DMSO treated CCLF_PEDS1012T, CCLF_PEDS_0022_T, and AFLAC_2377 cells were subjected to cell cycle analysis following flow cytometry. Bar graph representing the proportion of cells across phases of the cell cycle in at least biological replicates and 50k cells counted. **(d)** Deletion of *TP53* significantly increased the IC_50_ of KPT-330 in CCLF_PEDS1012T and CCLF_PEDS_0022_T. The fold change is based on comparison to a LacZ non-targeting control and based on biological duplicates. Error bars represent mean ±SD. **(e)** Change in viability using shTRIP13_1 and shTRIP13_2 across FHWT cell lines as compared to shControl and seed control (to shTRIP13_2). **(f)** Volcano plot showing the distribution of significant genes up or downregulated following sh*TRIP13*. RNA seq was performed on Aflac_2377T and CCLF_PEDS_0041_T cell lines with sh*TRIP13* as compared to shControl. Biological replicates performed. *TRIP13* is downregulated along with *CCND1*. **(g)** Gene set enrichment analysis (GSEA) enrichment score curves for the RB1/RBL1 skin specific knockout and G2M pathways following suppression with sh*TRIP13*. **(h)** Commonly upregulated and downregulated genes seen in both KPT-330 treated or sh*TRIP13* treated cells. Eight genes were upregulated and four genes were downregulated.

We found gene sets affecting both G1 (e.g. E2F) and G2/M pathways (**Fig 3b**) along with activation of the TP53 pathway (**Supp Table 2**). Other processes such as apoptosis were also enriched. We performed fluorescence-activated single cell analyses to assess the changes in the cell cycle following KPT-330 treatment. We found that changes in G1 were not consistent across our FHWT cell lines (**Fig 3c**). However, we saw decreases of S phase and consistent increases in G2/M suggesting that KPT-330 in our FHWT cell lines led primarily to a G2/M arrest (**Fig 3c**). We then confirmed the activation of TP53 through nuclear accumulation of TP53 following treatment with KPT-330 by 48 hours (**Fig 2e; Supp fig 3a**). As KPT-330 has been associated with upregulating nuclear accumulation of TP53, we sought to determine if TP53 was indeed an important mechanism within WT ^40^. We depleted *TP53* using CRISPR-Cas9 in PEDS1012T and PEDS_0022_T as compared to a control gRNA to *LacZ* (**Supp Fig 3b**). We then assessed the IC_50_ of KPT-330 and found a 6 to 13-fold increase in these values when *TP53* was deleted (**Fig 3d, Supp Fig 3c**). This significant change in IC_50_s suggests a critical role of *TP53* in KPT-330 induced cell death in FHWT.

We subsequently assessed the 412 differentially expressed genes and observed that the expression of *TRIP13*, Thyroid Hormone Receptor Interactor 13, was significantly downregulated in the KPT-330 treated cells (**Fig 3a**). TRIP13 previously was found to interact with co-factors of TP53 in injured renal epithelial cells ^41^. More recently, TRIP13 has been identified as a cancer predisposition gene in WT ^42^.

We then focused on the function of TRIP13 in our FHWT cell lines. First, we observed that in general, pediatric renal tumors had modest increases in expression of *TRIP13* (**Supp Fig 3d**) as compared to adjacent normal kidney controls. We then suppressed the expression of *TRIP13* with RNAi in *TP53* wild-type WT cells using two different *TRIP13* shRNA constructs (**Methods**). To mitigate off-target effects, we used a seed control to one shRNA construct and an RFP non-targeting control ^43^. Our cell viability results showed significant cell death in our cell lines transduced with either *TRIP13* shRNA constructs (14-59% cell viability; **Fig 3e and Supp Fig 3e–f**). In addition, when we overexpressed TRIP13 in our FHWT cell lines, we found a modest 15-27% increase in cell counts as compared to a luciferase overexpression control (**Supp Fig 3g**). We then tried to rescue our shTRIP13 viability defect using this overexpression vector but the 3’ UTR shRNA alone did not lead to a viability defect or suppression of TRIP13 by immunoblot (**Supp Fig 3e; shTRIP13-3**). However, in our TP53 deleted cells, we found that TRIP13 levels elevated at baseline and did not decrease upon KPT-330 treatment (**Supp Fig 3h**). These results suggest that FHWT (e.g., TP53 wild type) cells require TRIP13 for survival and when overexpressed, lead to a modest increase in proliferation.

We then assessed the transcriptional changes by RNA sequencing seen when we suppress *TRIP13*. We used the CCLF_PEDS_0041_T1 and AFLAC_2377 cell lines to determine the consequences of suppressing *TRIP13* as compared to our non-targeting shRNA controls (**Methods**). We found 172 genes upregulated and 101 genes downregulated (**Fig 3f**). We used GSEA to identify gene sets significantly enriched upon suppression of *TRIP13*. We found the gene sets involving RB1/RBL1 skin specific knockout mice as significant which was confirmed with decreased cyclin D1 levels (**Fig 3f–g; Supp fig 3f**) ^44^. We further found, similar to XPO1 inhibition, that TRIP13 is anti-correlated with the G2/M checkpoint (**Fig 3g**).

We subsequently asked how suppression of TRIP13 contributed to the phenotypes seen when FHWT cells were treated with KPT-330. We looked at the overlap between differentially expressed genes from our RNA-sequencing experiment (**Fig 3h**). We found eight genes upregulated and four genes downregulated. Interestingly, upregulated genes *CDH1*, *MYCT1* and *SAMD9* have been implicated as tumor suppressors in other cancers such as hematologic malignancies^45–47^. When we evaluated the down-regulated genes with TRIP13 suppression, we find genes involved in the cell cycle such as *ASPM* and *CDCA7*. ASPM is essential for mitotic spindle function in neurons ^48^ and CDCA7 regulates CCNA2, a cyclin with roles in G1 and G2/M, in esophageal squamous cell carcinomas ^49^.

In sum, our findings show that treatment with KPT-330 in part leads to suppression of TRIP13 which in turn leads to alterations in the cell cycle.

### KPT-330 and doxorubicin are synergistic *in vitro* and *in vivo* in WT

Doxorubicin was added to vincristine and dactinomycin in Stage III FHWT patients in the 1980s to account for higher risk disease (e.g., pulmonary metastasis or tumor rupture) ^50^. Despite improved overall survival with the addition of doxorubicin in patients with higher risk FHWT, these cancers recur in 15% of patients ^8^. Over the past two decades, only one Phase II clinical trial for relapsed FHWT has been opened (clinicaltrials.gov: NCT04322318). This trial uses a chemotherapy backbone without novel therapeutic targets. Thus, there is a need to identify potential synergistic combination strategies which could be tested in the Phase I/II setting.

We assessed the role of adding KPT-330 to doxorubicin to determine if there was synergy with our cell line models of FHWT. We first evaluated doxorubicin sensitivity in these patient-derived WT cell lines. We found that all WT were sensitive to doxorubicin with IC_50_s in the nanomolar range (~20-500nM) as compared to the normal cell lines which had an IC_50_ range of 0.65-1.5uM (**Fig 4a**).

**Figure 4:**
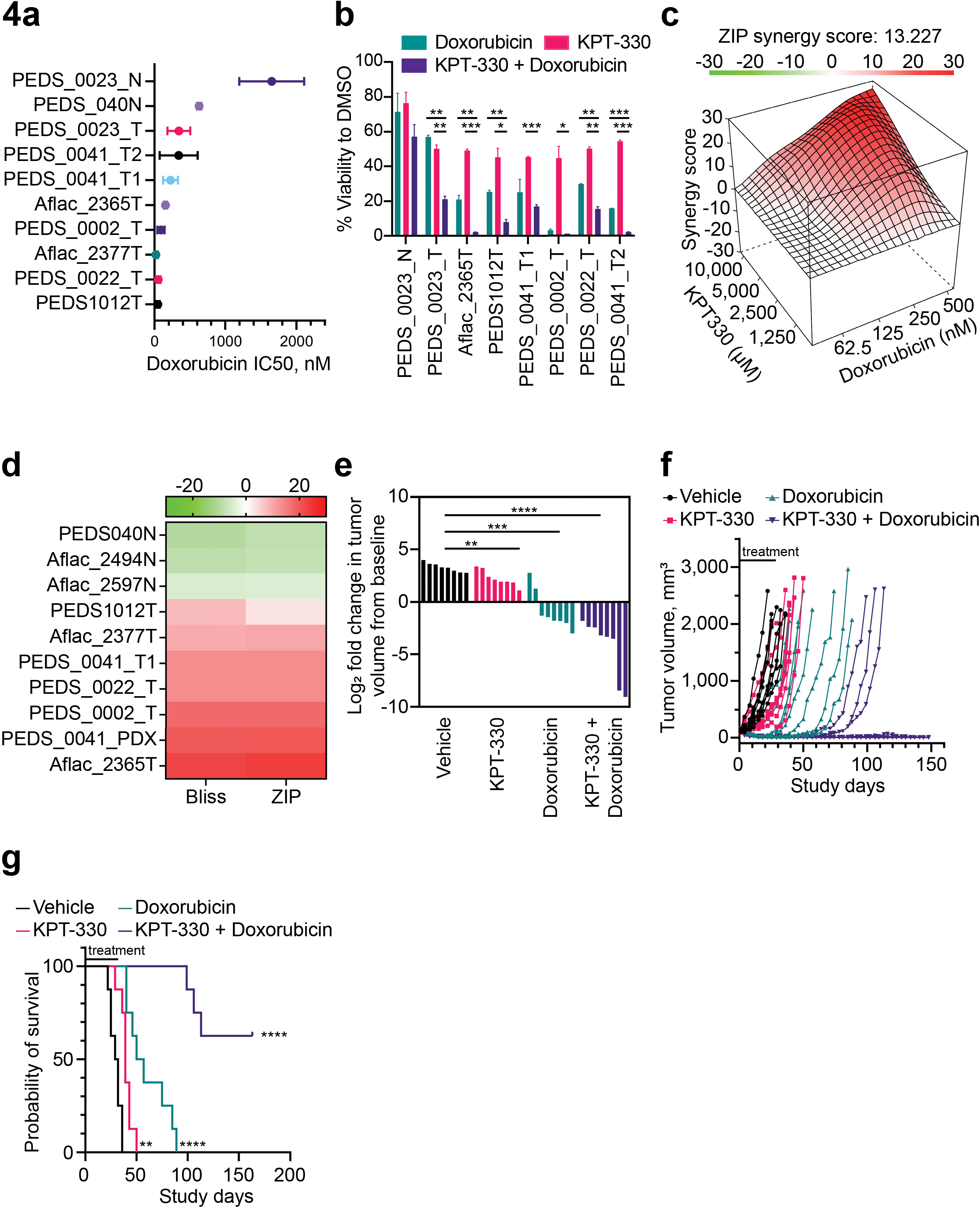
Combination of doxorubicin and KPT-330 is synergistic *in vitro* and *in vivo*. **(a)** IC_50_s for WT cell lines treated with doxorubicin as compared to normal kidney cells. X-axis is nM. SD shown from at least 2 biological replicates. **(b)** Bar graph representing the percent viability of WT cells upon treatment with doxorubicin (100nM) + KPT-330 (5μM) for 72 hours. Error bars represent mean ± SD from biological triplicates. **(c)** Representative 3D landscape image of the synergy score for WT cells treated with doxorubicin (62.5 - 500nM) and KPT-330 (1.25 – 10 μM). **(d)** Heat map showing synergy scores from Bliss and ZIP drug-drug interaction models in WT and normal kidney cells. Range green (antagonistic) to red (additive 0-10 to synergy >10). **(e)** CCLF_PEDS_0041 fragments were placed subcutaneously into the hind flank of NSG (NOD-SCID IL2Rgamma null) female mice. The tumor xenografts were treated with vehicle, doxorubicin, KPT-330 or the combination of doxorubicin and KPT-330 for 28 days. Log fold change of tumor volumes were calculated as compared to time of treatment (average 114.3 mm^3^). ** p-value <0.005, *** p-value <0.0005, **** p-value<0.00005. **(f)** Line graph depicting the tumor volume across the study days for each mouse in different treatment groups. Treatment lasted 4 weeks from study enrollment and continued until mouse met endpoint or study termination at day 150. **(g)** Kaplan Meier survival curves representing the probability of overall survival in the patient-derived xenografts (n=8 per treatment group) treated with the vehicle, KPT-330, doxorubicin and KPT-330 + doxorubucin. ** p-value <0.005, **** p-value<0.00005.

We then evaluated the effect of KPT-330 and doxorubicin treatment in combination on cell viability as measured by CellTiter-Glo. The addition of KPT-330 to doxorubicin significantly enhanced cell death in WT cells (**Fig 4b**). We then performed synergy experiments (**Methods**) to determine if this combination was synergistic or additive. Across multiple concentrations of KPT-330 and doxorubicin in all of the WT cell lines tested, the combination was synergistic (e.g., scores >10; **Fig 4c–d**). In contrast, this combination was antagonistic in normal cells (**Fig 4d**). Collectively, we found a synergistic interaction between KPT-330 and doxorubicin in FHWT as compared to normal kidney cell lines.

We followed up these *in vitro* studies with *in vivo* studies to further validate the potential synergistic effect of KPT-330 and doxorubicin. Of our patients samples, we were able to generate two PDXs from CCLF_PEDS_0041 and CCLF_PEDS_0023. Given that CCLF_PEDS_0023 has features of anaplastic Wilms, we performed our *in vivo* studies with the PDX from CCLF_PEDS_0041, a patient with FHWT. Interestingly, the CCLF_PEDS_0041_T1 cell line and cell line derived from the PDX (CCLF_PEDS_0041_T2) were some of the least sensitive WT cell lines to KPT-330 and doxorubicin treatments (**Fig 2d and 4a–b**). We treated tumor xenografts with placebo, doxorubicin, KPT-330 or the combination of doxorubicin and KPT-330 for 28 days and then monitored tumor growth until endpoints were reached or after 150 days following treatment initiation. We found at 22 days, that KPT-330 led to 49% decrease (p-value = 0.046), doxorubicin led to 87% decrease (p-value < 0.005) and the combination led to 99% decrease (p-value < 0.005) in tumor volume compared to the vehicle (**Fig 4e**). We then assessed the effects following cessation of therapeutic treatment. We found that monotherapy with KPT-330 or doxorubicin led to a median survival of 39 and 53.5 days, respectively (**Fig 4f–g**). Further, the combination of KPT-330 and doxorubicin had an undefined median survival with five of eight mice exhibiting a complete response (**Fig 4f–g**). Taken together, these observations suggest that inhibition of XPO1 in combination with doxorubicin chemotherapy are synergistic *in vivo*.

## DISCUSSION

Here we have developed short term cell line models of Wilms Tumor (**Fig 1**). Although these cell lines, which maintain a faithful representation of their genomics and transcriptomics, only grow for a finite number of passages, we show that they can be used to perform systematic loss-of-function studies (**Fig 2**). We find that integration of targeted RNA interference and CRISPR-Cas9 screens identifies the nuclear export apparatus as a therapeutic target in FHWT.

XPO1 regulates a spectrum of cellular processes by controlling the active transport of over 200 proteins out of the nucleus, including tumor suppressors (TSs) and transcription factors (TFs) (such as TOP2A, p53, WTX, APC, Rb, and PI3K/AKT) ^3^. Due to its important physiological role, inhibition of XPO1 by the small molecule inhibitor, KPT-330 (selinexor), is widely being tested in the phase I and phase II clinical trials in multiple malignancies and has FDA approval in several hematologic malignancies ^4–6^. As part of the Pediatric Preclinical Testing Program, KPT-330 was tested as a single agent across a panel of pediatric hematologic and solid tumor PDXs which included several WT xenografts ^51^. In this study, the group used KPT-330 in 3 WT PDXs. They found one PDX (KT-10) had maintained a complete response at 42 days post-treatment initiation while the other two had partial responses. However, mechanisms and biomarkers remain unknown.

Here, we show that inhibition of nuclear export leads to primarily a G2/M arrest in FHWT and loss of TP53 leads to significant resistance to KPT-330 (**Fig 3)**. While the role of TP53 in response to KPT-330 has been previously reported^52^, we identified that suppression or inhibition of nuclear export leads to suppression of TRIP13 in FHWT. We find that although TRIP13 is traditionally thought to affect G2/M^53,54^, loss of TRIP13 affects both G1 and G2/M in FHWT. Furthermore, when we compare the differentially expressed genes between KPT-330 treated cells as compared to cells with *TRIP13* suppression, we find several potential tumor suppressors (e.g. *CDH1*, *MYCT1*) and *SAMD9*, a gene having antiproliferative properties, whose expression increases. Future studies will explore these findings to better elucidate the function of TRIP13 in FHWT. Finally, we show that there is significant synergy when KPT-330 is combined with the topoisomerase II inhibitor, doxorubicin, *in vivo* where five of eight mice achieved cure following a single cycle of combination therapy (**Fig 4**).

Future work requires the development of additional 1q gain or 1p/16q loss of heterozygosity (LOH) *in vitro* models. In addition, our data from one patient derived model with DAWT (TP53 mutant) suggests that inhibition of nuclear export may be a therapeutic strategy but additional models and validation studies are needed (**Fig 2b**).

Taken together, we have identified that KPT-330 acts in part by inhibiting TRIP13 in Wilms Tumor. In addition, suppression of *TRIP13* may serve as a biomarker of response in WT. Finally, in combination with a topoisomerase II inhibitor, nuclear export inhibitors could be studied in patients with high-risk Favorable Histology WT.

## METHODS

### Development of patient-derived Wilms tumor cell lines

Tumor cells were isolated from tumor and if available, adjacent normal kidneys tissues of patients with a pathological diagnosis of Wilms Tumor. Samples were obtained under IRB approved protocols from patients at either Dana Farber / Boston Children’s Cancer and Blood Disorders Center or Aflac Cancer and Blood Disorders Center at the Children’s Healthcare of Atlanta and Emory University School of Medicine. Tumor tissue or adjacent normal tissue was first minced into 1-2mm^3^ pieces and resuspended in F-media as previously described^23^. Samples were then plated in six-well plates. Cells were serially passaged after reaching confluency of 70-80%. For PDX cell lines, a tumor fragment (<5mm in diameter) was implanted into NSG mice and once passaged twice, a tumor fragment was then minced and fragmented on a 6 well dish to develop a PDX cell line. In general Wilms Tumor cells were grown for up to 30 passages and normal kidney cells were grown for approximately 12-15 passages before senescing.

### Low coverage whole genome sequencing (WGS) and Copy Number Analyses

DNA from tumor tissue, normal tissue if available or blood, and cell pellets from our patient-derived cell lines was extracted (NEB, T3010S). Genomic libraries were prepared (Illumina) by Novogene Inc and sequenced at 0.1x on Novoseq 6000 (Illumina). Fastq or BAM files were mapped and aligned using Illumina Dragen v3.7.5 to GrCh38 on Amazon Web Services. ichorCNA v0.10 was then used to determine copy number alterations ^55^. Copy number variations (amplification, gain, deletion) were calculated by ichorCNA.

### Whole exome sequencing

WES was performed using DNA as described above. For five patients, WES was performed at the Broad Institute Genomics Platform using HiSeq 2000 (Illumina). WES libraries were based on an Illumina Customized Exon (ICE) array. For three patients, WES was performed at Novogene Inc using Novoseq6000 (Illumina). Libraries were based on an Agilent array. Sample coverage was >100x for tumor and >50x for normal. Fastq or BAM files were mapped and aligned using Illumina Dragen v3.7.5 to GrCh38^56^ on Amazon Web Services. Variant call files (vcf) were subsequently processed with OpenCRAVAT using cancer hostpots, cancer gene landscape, chasm plus, civic, cosmic, mutpanning and ndex NCI to filter for pathogenic variants.

### RNA sequencing

RNA was extracted from samples collected from tumor, adjacent normal and cell lines (Qiagen RNeasy or NEB Monarch Total RNA). Libraries were prepared using Illumina TruSeq. Samples were run with at least 50 million paired-end reads using Novoseq 6000 (Illumina). For mechanistic studies, RNA was collected in biological replicates from cells treated with shRNAs or compounds as listed in the figures. Similar methods were used in sequencing with the exception that samples were sequenced at 20 million single-end reads per sample. Fastq or BAM files were mapped and aligned using Illumina Dragen v3.7.5 to GrCh38^56^ and GenCode36^57^ on Amazon Web Services. Samples were then quantified using salmon through Illumina Dragen v3.7.5. The quantification files were imported into R using tximport ^58^ and ComBat-seq ^59^ was used to correct for batch effects. Differential gene expression analysis was then performed using DESeq2 ^60^. UMAP ^61^ was used for visualizing clustering between samples.

### Loss of function screens – methodology and analyses

RNAi and CRISPR-Cas9 screens were performed as previously described ^21–23^. Specifically, we transduced into noted cell lines the DCT v1.0 libraries: shRNA (CP1050) and sgRNA (CP0026) libraries from the Broad Institute Genetic Perturbation Platform (GPP) (http://www.broadinstitute.org/rnai/public/). In parallel, CCLF_PEDS_0023_T, CCLF_PEDS_0023_T2, and CCLF_PEDS_0002_T cells were transduced with Cas9 expression vector pLX311_Cas9^62^. These stable cells were subsequently transduced using the DCT v2.0 libraries and at a multiplicity of infection (MOI) between 0.3-0.6^32^. Screens were performed goal representation rate >500 cells per shRNA or sgRNA. Cells were passaged every 5-7 days until days 25-29 following transduction. Genomic DNA was extracted from an early time point and at the end time point. Samples were sequenced as previously described, deconvoluted and analyzed using MAGeK^33^.

### Cell viability assays

Cells were plated in a concentration of 2,500 cells/well in a 96 well plate (Corning, 3903, or Greiner Bio-One, 655098) and were incubated overnight in F-media overnight. After 24 hours, F-media was aspirated and F-media with the indicated drug concentration was added. Cells were treated between range of 0.007 μM to 50 μM of KPT-330 (Selleck, S7252) and doxorubicin (Selleck, S1208) for 72 hours. Cell viability was measured using CellTiter-Glo (Promega, G7573). IC_50_s of KPT-330 and doxorubicin were determined in each cell line by fitting the dose response curve using Graphpad Prism v9.3.0. Each experiment was repeated in at least two biological replicates.

### XPO1 and TRIP13 shRNA experiments

shRNA sequences were designed using the Broad Institute GPP portal (http://www.broadinstitute.org/rnai/public/). Oligos were obtained from Integrated DNA Technologies (IDT). Oligos were annealed and ligated with pLKO.5 as previously described. Constructs were transfected using TransIT-LT1 (Mirus Bio LLC, Madison, WI, USA) and HEK 293T cells. Cells were then transduced with lentivirus to achieve appropriate knockdown without significant viral toxicity as previously described ^22^. Following selection with puromycin (Invivogen), cell proliferation was assayed by CellTiter-Glo at 10 days. Experiments were repeated a minimum of three times in technical triplicates. shRNA sequences can be found in **Supp Table 4**.

### *TP53* CRISPR-Cas9 deletion

CCLF_PEDS1012T and CCLF_PEDS_0022_T cells were transduced with Cas9 expression vector pLX311_Cas9. These cells were subsequently transduced with sgTP53-1 and sgLacZ in the pXPR003 backbone. Transduced cells were then cultured in the indicated antibiotics for selection and suppression of gene expression was confirmed by immunoblotting.

### qRT-PCR

Total RNA was extracted using the NEB Monarch RNA extraction kit. One microgram of RNA and oligo primers were used for cDNA synthesis in a total reaction volume of 20 µL High-Capacity cDNA Reverse Transcription Kit (Thermo Fisher Scientific, 4368814). qRT-PCR reactions were prepared using SYBR-Green PCR Master Mix (ThermoFisher Scientific, 4367659) and run on a BioRad CFX96 qPCR System with a minimum of technical duplicates. Relative mRNA levels were calculated using the 2-ΔΔCt method^63^. Sequences of primers used in qRT-PCR experiments are detailed in **Supp Table 3**. Results shown are representative of at least two biological replicates.

### Immunoblots

Cells were grown to 70-80% confluency and were treated or transduced as indicated the figures. Thereafter, cells were lysed using 1x RIPA (Cell Signaling Technologies, 9806) with protease inhibitors (coMplete, Roche, 42484600) and phosphatase inhibitors (PhosSTOP, Roche, 04906837001). For experiments using nuclear and cytoplasmic fractions, fractions were extracted using NE-PER (ThermoFisher Scientific, 78835). Using 10% or 4-12% SDS-PAGE gels, gels were transferred onto PVDF (Millipore, IPFL00010) or nitrocellulose membranes (ThermoFisher Scientific, IB23001). Blots were then visualized using Odyssey CLx (Licor, Lincoln, NE). Antibodies used in this study include: XPO1 (Santa Cruz; sc-5595), β-Actin (C-4) (Santa Cruz; sc-47778), β-Actin (Cell Signaling; 8457), TP53 (Santa Cruz; sc-123), TRIP13 (Abcam; ab128171), α-Tubulin (Santa Cruz, sc-5286), α-Tubulin (Cell Signaling; 2144), Lamin A/C (Cell Signaling; 4777 or 2032), p21 (Cell Signaling; 29475), CCND1 (Santa Cruz; sc-8396). Results shown are representative of at least two biological replicates.

### Synergy Experiment

WT cells were plated in 96 well plates at a concentration of 2,500 cells/well. After 24 hours, cells were treated with a combination of KPT330 and doxorubicin in a 5×5 matrix on the same microtiter plate in duplicate. Each plate included a 4-dose dilution of each drug alone or in combination and included DMSO controls. Following a 3-day incubation, we performed a cell viability assay using CellTiter-Glo. Relative cell viabilities were then calculated and analyzed with SynergyFinder^64^ using two alternate drug-drug interaction models: zero interaction potency (ZIP)^65^ and bliss independence^66^. Experiments were performed in biological replicates or triplicates.

### *In vivo* experiments

Female NSG (NOD-scid IL2Rgamma null) mice, 6-weeks old were purchased from The Jackson Laboratory (Bar Harbor, ME). Animals were acclimated for at least 5 days before initiation of the study. The study was conducted at Dana-Farber Cancer Institute (DFCI) with the approval of the Institutional Animal Care and Use Committee in an AAALAC accredited vivarium.

NSG mice were implanted with CCLF_PEDS_0041_T2 tumor fragments from previously expanded tumors, subcutaneously in the hind-flank. Tumors were allowed to establish to 86 – 163 mm3 (average 114.3 mm3) in size before randomization using Studylog software (San Francisco, CA) into various treatment groups with 8 mice/group as follows: vehicle control (0.6% Pluronic F-68 + 0.6% Plasdone PVP; oral gavage on MWF x 4 weeks), selinexor (purchased from Selleck Chemicals LLC, 15 mg/kg oral gavage on MWF x 4 weeks), doxorubicin (obtained from DFCI pharmacy, 5 mg/kg intravenous injection on days 2 and 9) and the combination of selinexor and doxorubicin. No statistical methods were used to pre-determine sample sizes but our sample sizes are similar to those reported in previous publications ^22,23,67^. Once the treatment was completed, tumors were monitored at least once a week. For single agent and combination agent efficacy studies, mice were dosed for 21 or 28 days and monitored daily. Drugs were then withdrawn, and tumors were monitored twice weekly until study termination on day 150. Tumor volumes were calculated using the following formula: (mm3) = length × width × width × 0.5. Mice were immediately euthanized if the tumor volume exceeded 2000 mm3 or if the tumors became necrotic or ulcerated. The compounds were well tolerated with less than 8% body weight loss. Data collection and analysis were not performed blind to the conditions of the experiments. GraphPad Prism v9.3.0 was used to calculate significance in the Kaplan-Meier curves.

## AVAILABILITY OF DATA AND RESOURCES

Sequencing data reported in this paper has been deposited in the database _PENDING_.

## COMPETING INTERESTS

The authors declare the following competing financial interests – K.M. is currently employed at PreludeDx. A.L.C. is currently an employee of Aztra Zeneca. G.C. is currently an employee of Johnson & Johnson. D.M.W is currently an employee of Merck and has research support from Daiichi Sankyo, Abcuro, Verastem, Secura, and is on the Advisory Board or has equity in Ajax, Travera, Astra Zeneca, Bantam. K.S. receives grant funding from Novartis and KronosBio, consults for and has stock options in Auron Therapeutics and has served as an advisor for KronosBio and AstraZeneca. W.C.H. is a consultant for Thermo Fisher, Solasta Ventures, MPM Capital, KSQ Therapeutics, iTeos, Tyra Biosciences, Jubilant Therapeutics, RAPPTA Therapeutics, Function Oncology, Frontier Medicines and Calyx. The remaining authors declare that they have no competing interests.

## FUNDING

Research reported in this publication was supported by the following: NIGMS T32GM008490 (GWC), T32GM007739 (WJK), NCI K08 CA2555569-01 (AJM), NCI R35 CA231958 (DMW), NIH R35 CA210030 (KS), NCI U01 CA176058 (WCH), ACS MRSG-18-202-01 (ALH). CureSearch for Children’s Cancer YIA (ALH), Rally for Childhood Cancer Investigator Grant 21IN12 (ALH).

## AUTHOR CONTRIBUTIONS

KM, WCH and ALH in the design of the study. KM, BPL, WCH and ALH wrote the manuscript. KM, BPL, GWC, BD, MR, MB and ALH performed analyses of the sequencing data. KM, BPL, WJK, MD, BK, PK, MT, JB, ALH generated the cancer cell lines. JS, HCJ, TL and KCG oversaw the Aflac Solid Tumor Biorepository. SM, CC, and BC oversaw the DF/BCH Solid Tumor Biorepository. WJK, MD, DA, MK, XY, GC, DR and ALH performed and/or analyzed the functional genomic screens. ALC, BL, CM, AJM, KL, DMW, PG generated the PDXs and performed in vivo studies. MB, BC, MT, JB, KL, DR, AJM, DMW, PG, JS, MR, EAM, KS, KCG, WCH and ALH supervised the studies. All authors discussed the results and implications and edited the manuscript. All authors also read and approved the final manuscript.

## ACKNOWLEDGEMENTS

We thank the patients and their families for their participation. We thank the Hong, Spangle and Hahn labs for their thoughtful comments and suggestions. Tissue samples were provided by the Children’s Healthcare of Atlanta Pediatric Bio-Repository or Dana-Farber/Boston Children’s Center for Cancer and Blood Disorders. Other investigators may have received specimens from the same subjects. Research reported in this publication was supported in part by the Pediatrics/Winship Flow Cytometry Core of Winship Cancer Institute of Emory University, Children’s Healthcare of Atlanta and NIH/NCI under award number P30CA138292, by the Bioinformatics and Systems Biology Shared and the Biostatistics Shared Resource of Winship Cancer Institute of Emory University and NIH/NCI under award number P30CA138292. The content is solely the responsibility of the authors and does not necessarily represent the official views of the National Institutes of Health.

## SUPPLEMENTARY FIGURES AND SUPPLEMENTARY FIGURE LEGENDS

**Supp Fig 1:**
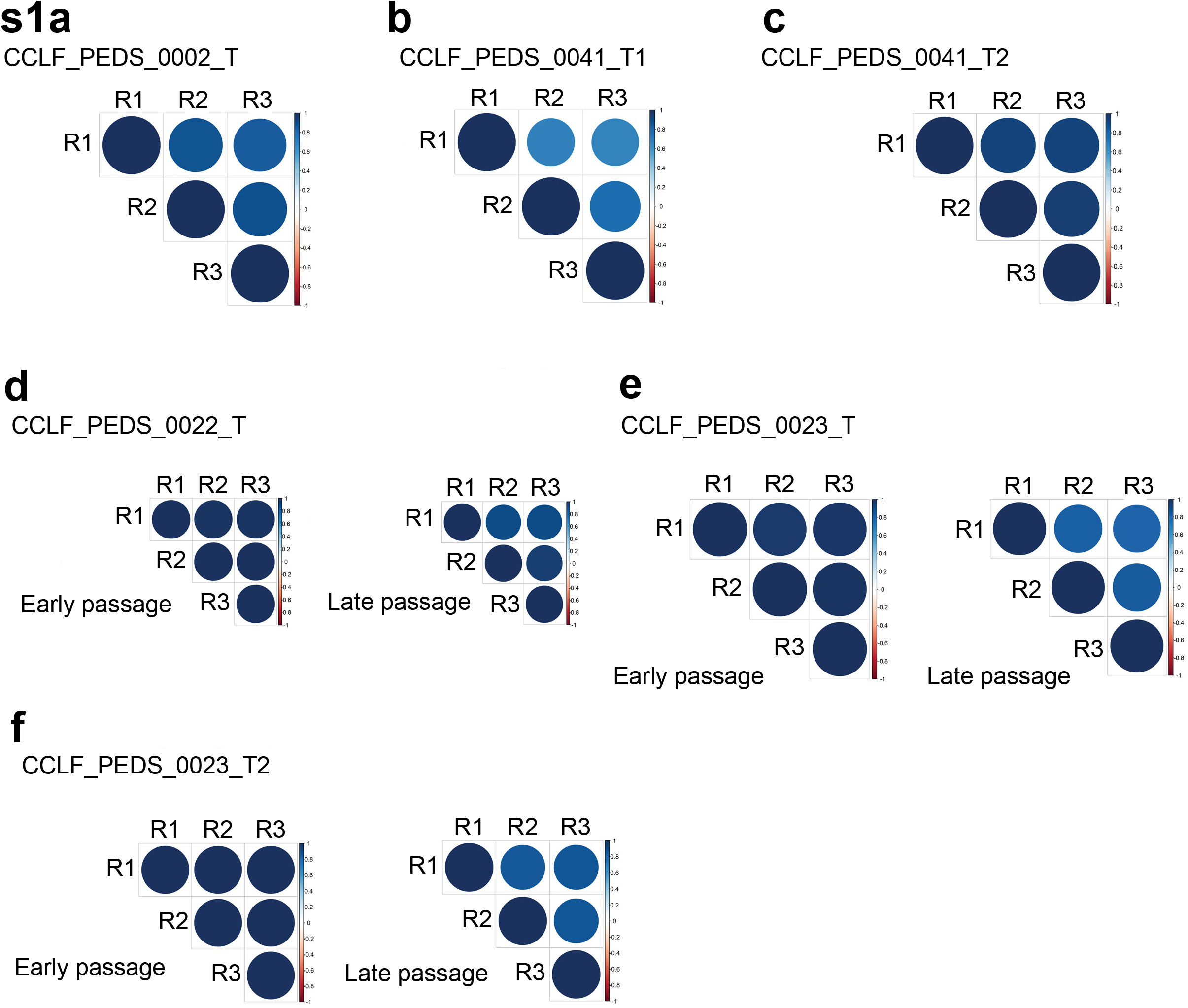
Tight correlation between shRNA and sgRNA loss of function screens among biological replicates. RNAi screens represented in a-c. **(a)** CCLF_PEDS_0002_T, **(b)** CCLF_PEDS_0041_T1, **(c)** CCLF_PEDS_0041_T2. CRISPR-Cas9 screens used early and late passages to determine change in abundance of sgRNAs as shown in d-f. **(d)** CCLF_PEDS_0022_T, **(e)** CCLF_PEDS_0023_T, **(f)** CCLF_PEDS_0023_T2. Right color gradient delineates the Pearson’s correlation with anticorrelation as red and correlation as blue.

**Supplementary Figure 2:**
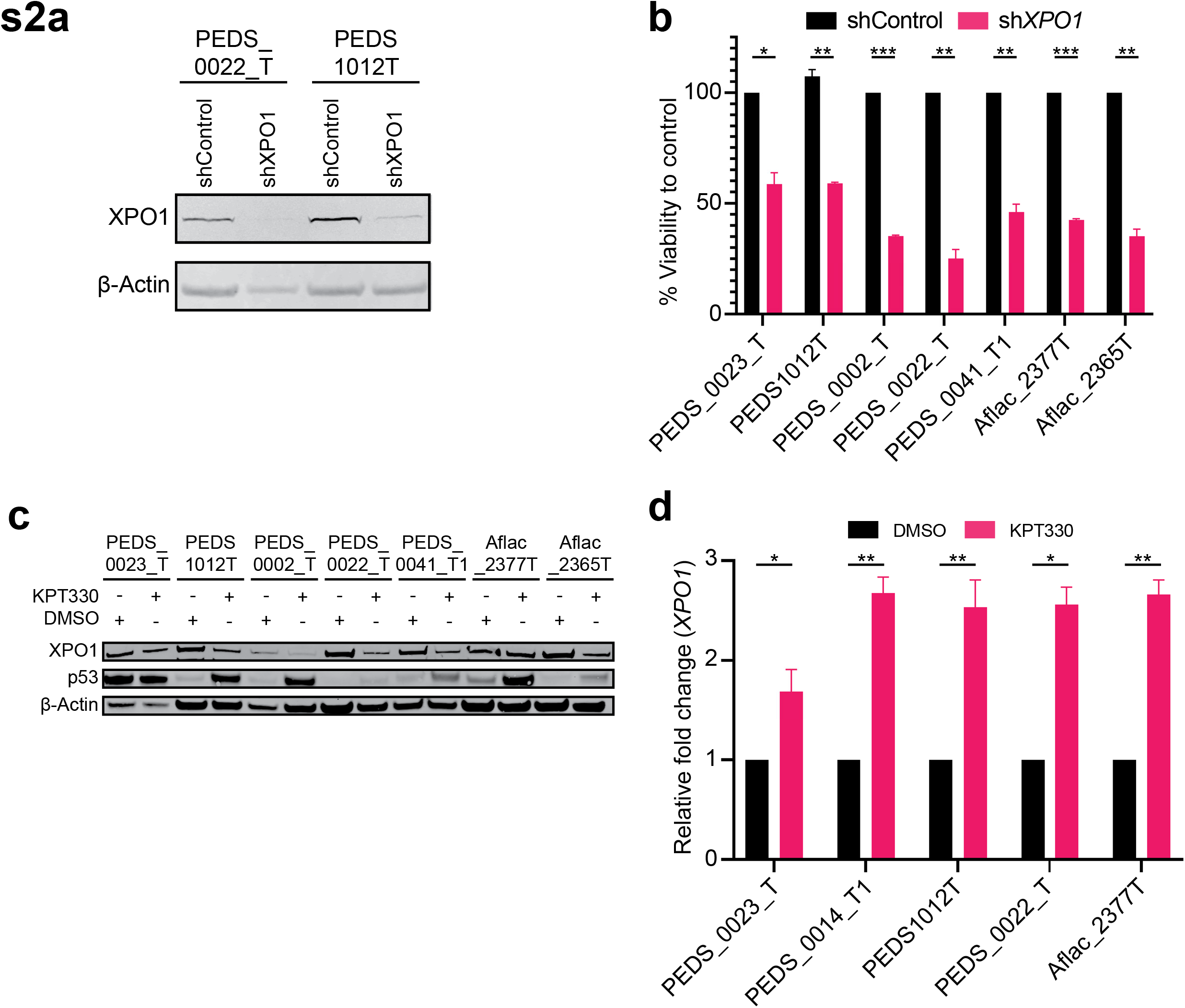
Inhibition of nuclear export in FHWT. **(a)** Suppression of XPO1 by immunoblot **(b)** Change in viability using sh*XPO1* across FHWT cell lines as compared to shControl **(c)** At 24 hours, XPO1/CRM levels are suppressed with accumulation of TP53 across a majority of cell lines. **(d)** qRT-PCR analysis of *XPO1* in KPT-330 and DMSO treated cells. Data was normalized by the amount of TBP, expressed relative to the corresponding value for all the cells and are means ± SD from at least two biological replicates. * indicates a p-value<0.05, ** p-value <0.005, *** p-value <0.0005.

**Supplementary Figure 3:**
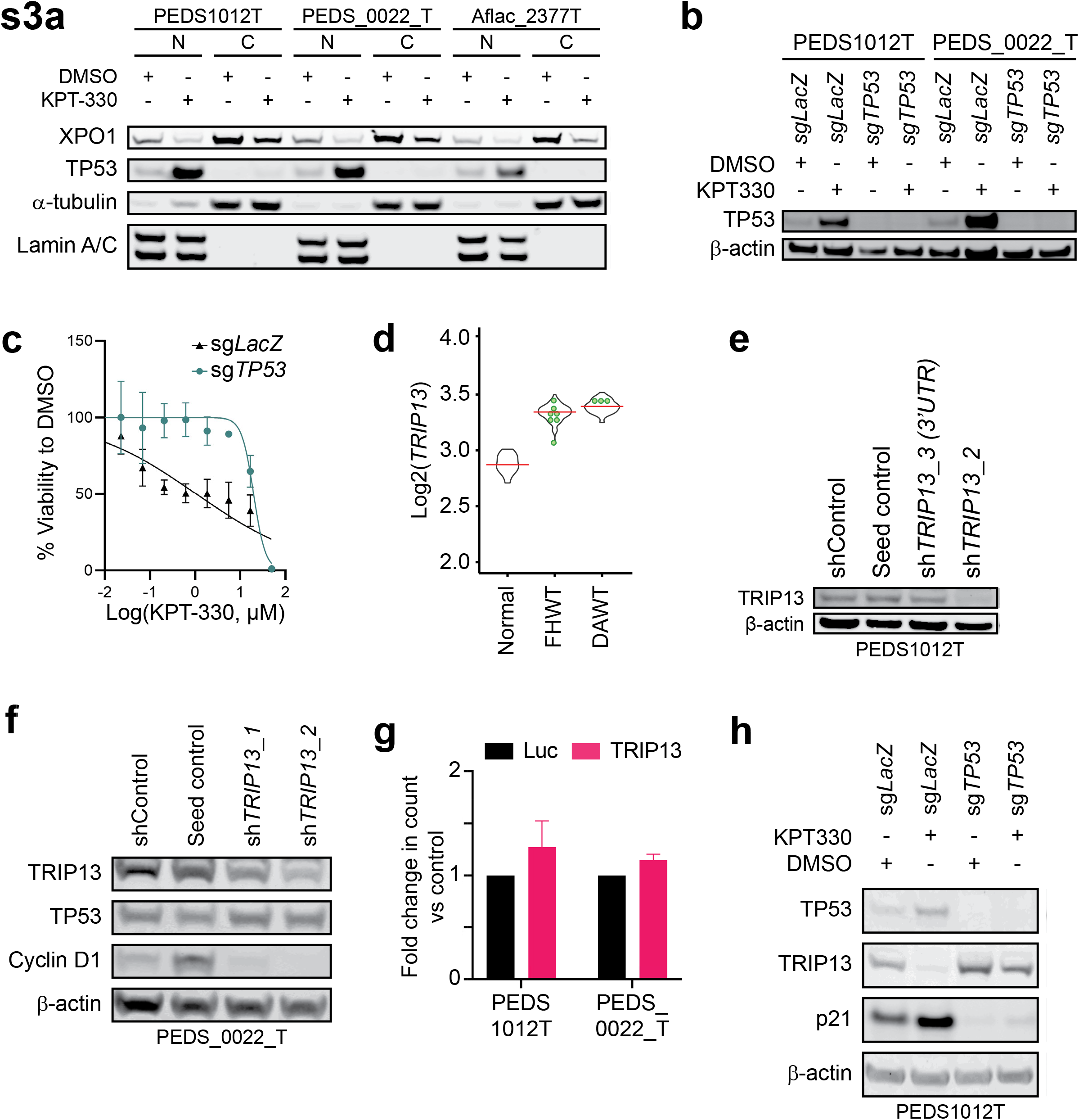
Inhibition of nuclear export and TRIP13 in FHWT cell lines. **(a)** Immunoblots depicting the decrease in protein levels of XPO1 in the cytoplasmic lysates and nuclear accumulation of TP53 in the nuclear lysates upon treatment with KPT-330 at 24 hr. **(b)** We introduced sgRNAs targeting either *LacZ* or *TP53* and confirmed decrease in TP53 following treatment with KPT-330. **(c)** Dose-response curves for the sg*TP53* and sg*LacZ* cells for KPT-330. Error bars represent mean ±SD and represent biological replicates. **(d)** Dot plots representing the expression levels of *TRIP13* in Wilms tumor when compared with the normal matched kidney tissue. Immunoblot depicting suppression of TRIP13 in **(e)** CCLF_PEDS1012T **(f)** CCLF_PEDS_0022_T **(g)** Overexpression of TRIP13 leads to modest increase in proliferation as compared to luciferase. **(h)** TP53 and p21 protein levels increase while TRIP13 levels decrease upon KPT-330 treatment in TP53 wildtype cells.

